# A high sensitivity strategy to screen NAD(P)H-dependent oxidoreductase activity by coupled enzyme cascade

**DOI:** 10.1101/2025.06.15.659766

**Authors:** Trisha Ghosh, Jacob Sicheri, David H. Kwan

**Affiliations:** Department of Biology, Concordia University, 7141 Sherbrooke Street West, Montreal, Quebec, Canada, H4B 1R6; Department of Chemistry and Biochemistry, Concordia University, 7141 Sherbrooke Street West, Montreal, Quebec, Canada, H4B 1R6

## Abstract

Enzymes play a pivotal role in “green chemistry” as tools for biocatalysis. Oxidoreductase enzymes are especially useful for carrying out key electron transfer (redox) steps towards a wide range of chemical transformations (e.g., asymmetric hydrogenation, oxygenation, hydroxylation, epoxidation, or Baeyer-Villiger oxidation) that might not otherwise be available to chemists through conventional (nonbiological) synthetic approaches. The ability to screen oxidoreductase activity is important in identifying useful biocatalysts from nature, and also towards engineering novel ones through directed evolution. Many valuable redox enzymes are dependent upon NAD(P)H as an electron donating co-substrate (or conversely, upon NAD(P)^+^ as an electron acceptor), and the common method to detect their activity is to monitor the change in absorbance at 340 nm as NAD(P)H is converted to NAD(P)^+^ (or vice versa). The limited sensitivity of this method presents a challenge in detecting very low levels of oxidoreductase activity, and this can prove very difficult to begin engineering enzymes as improved biocatalysts when the rates of natural enzymes may be slow for a desired redox reaction. Herein, we report a fluorescence-based, enzyme cascade-coupled system that we have developed to detect oxidoreductase activity with orders of magnitude more sensitivity than conventional absorbance-based assays. While recycling NAD(P)H from NAD(P)^+^, the coupled enzyme cascade triggers cleavage of a fluorogenically labeled probe, releasing a strong fluorescent signal. This allows detection of very low levels of a specific oxidoreductase activity that we may wish to magnify by directed evolution using our assay in high-throughput screening.

## Introduction

Advances in enzyme engineering have enabled the development of new and improved biocatalysts for applications in green chemistry, bioprocessing, biotherapeutics, diagnostics, and pharmaceuticals [1, 2]. A key challenge in optimizing enzyme function through directed evolution is efficiently screening large variant libraries to identify beneficial mutations. This requires high-throughput assays capable of detecting enzyme activity and linking phenotype to genotype, ensuring that desired traits can be traced back to their genetic origins [3]. By enabling rapid and accurate evaluation of enzyme performance, these assays are essential for selecting and evolving variants with enhanced properties.

Strategies for detecting enzyme activity depend largely on the enzyme type and whether its function can be linked to a measurable output [4]. Coupled enzyme assays have been developed to enable detection of otherwise unmeasurable reactions by linking the activity of one enzyme to another that generates a quantifiable signal [5]. Ideally, such assays should be rapid, efficient, and high throughput. Similarly, multi-enzyme systems have been utilized for substrate recycling and cofactor regeneration, as well as for catalyzing reaction cascades that yield detectable outputs [6, 7]. These enzyme cascades can be designed to produce measurable signals, such as fluorescence or absorbance, facilitating enzyme activity screening. Beyond screening applications, enzyme cascades are also valuable for synthesizing high-value compounds in a sustainable manner [8–10]. By expanding the functional scope of biocatalysis, they enhance the applicability of individual enzymes or entire pathways. Developing high-throughput assays for these cascades is crucial for efficient enzyme screening and protein engineering. Furthermore, enzymatic cascades can be adapted for diverse screening and characterization purposes, making them a versatile platform. Thus, *in vitro* coupled enzyme assays serve as a powerful tool in cell-free enzyme engineering and directed evolution.

Oxidoreductases are commercially valuable enzymes that catalyze oxidation-reduction reactions, playing pivotal roles in various industrial applications including biofuels, food and agriculture, pharmaceutical and fine chemical synthesis, and even environmental bioremediation [10–12]. Examples include the use of enzymes like glucose and alcohol dehydrogenases in biofuel cells to convert chemical energy into electrical energy [13], production of raspberry ketone as a flavoring compound using benzalacetone reductase or P450 enzymes [14, 15], production of indigo dye for textiles by P450-catalyzed hydroxylation of indole [15, 16], Baeyer-Villiger monooxygenase-catalyzed synthesis of lactones as chemical precursors (*e.g.*, for drugs, and synthetic polymers) [17–19], and the use of ring-hydroxylating oxygenases to degrade polyaromatic hydrocarbons that are found as environmental pollutants [20].

Ene reductases, a subgroup of NAD(P)H-dependent oxidoreductases, exemplify the industrial potential of these enzymes through their roles in industrial biocatalysis [21]. Among them, Old Yellow Enzymes (OYEs), have garnered significant interest as biocatalysts due to their ability to catalyze the asymmetric hydrogenation of electron-deficient alkenes. Their broad substrate range allows them to act on a variety of electron-poor alkenes, including ketones, carboxylic acids, and aldehydes, making them valuable tools for green chemistry applications. Like many oxidoreductases, OYEs rely on NADH or NADPH as cofactors (as well as a noncovalently bound flavin mononucleotide (FMN) at their active site) [22, 23]. Their versatility and efficiency highlight the expanding role of oxidoreductases in sustainable industrial processes, particularly in fine chemical and polymer synthesis.

Given their widespread practical applications in biotechnology, optimizing oxidoreductases through enzyme engineering is of significant interest. Screening strategies for oxidoreductase catalytic activity primarily involve colorimetric detection and absorbance spectrometry. An important group of oxidoreductases utilizes nicotinamide adenine dinucleotide or nicotinamide adenine dinucleotide phosphate as cofactors, which can act as electron donors (NADH or NADPH) or electron acceptors (NAD⁺ or NADP⁺). The concentration of NAD(P)H can be determined by measuring absorbance at 340 nm, making it a convenient indicator for assessing the activity of NAD(P)H/NAD(P)⁺-dependent enzymes. While these methods can potentially be used to assay all NAD(P)H/NAD(P)^+^-dependent oxidoreductases, measuring a decrease in absorbance provides limited sensitivity that is insufficient to detect very low levels of activity especially in cases where background signal may be relatively high. Fluorescence based assays are generally preferred over absorbance due to their high sensitivity which help detect enzyme activity at low concentrations.

We developed a fluorescence-based, enzyme cascade-coupled system to detect oxidoreductase activity with vastly greater sensitivity than conventional absorbance-based assays. While recycling NAD(P)H from NAD(P)^+^, the coupled enzyme cascade catalyzes multiple transfers of a phosphate group between intermediates resulting in the phosphorylation of a fluorogenic probe, allowing its cleavage by a hydrolase that releases 4-methyllumbeliferone resulting in strong fluorescence. We thus established a continuous fluorescence-based assay to screen the activity of NAD(P)H-dependent redox enzymes with high sensitivity in a high throughput setup making it suitable for directed evolution.

## Results and Discussion

For the *in vitro* coupled enzyme cascade that we designed (**Fig. 1**), four reaction steps are linked downstream of an NAD(P)H-dependent redox reaction. The NAD(P)^+^ produced from the oxidoreductase-catalyzed reaction is used to trigger the cascade, which first involves a glyceraldehyde 3-phosphate dehydrogenase (GAPDH)-catalyzed reaction that converts glyceraldehyde-3-phosphate (G3P) and inorganic phosphate (P_i_) into 1,3-bisphosphoglycerate with the concomitant regeneration of NAD(P)H from NAD(P)^+^. The second step of the coupled cascade has phosphoglycerate kinase (PGK) catalyzing the conversion of ADP to ATP with a phosphate transferred from 1,3-bisphosphoglycerate, which is converted to 3-phosphoglycerate as a result. In the third step, a β-glucoside kinase, BglK, consumes ATP to phosphorylate 4-methylumbelliferyl β-D-glucoside (MU-Glc), producing 4-methylumbelliferyl 6-phospho-β-D-glucoside (MU-6P-Glc) and releasing ADP in the process. Fourth and finally, MU-6P-Glc is recognized by a 6-phospho-β-glucosidase, BglA-2, and hydrolyzed to release glucose-6-phosphate and fluorescent 4-methylumbelliferone. The enzymes used in the cascade—originating from *Escherichia coli* K-12 (GAPDH and PGK), *Klebsiella pneumoniae*, (BglK), and *Streptococcus pneumoniae* TIGR4 (BglA-2)—were recombinantly expressed in *E. coli* BL21(DE3).

**Figure 1.**
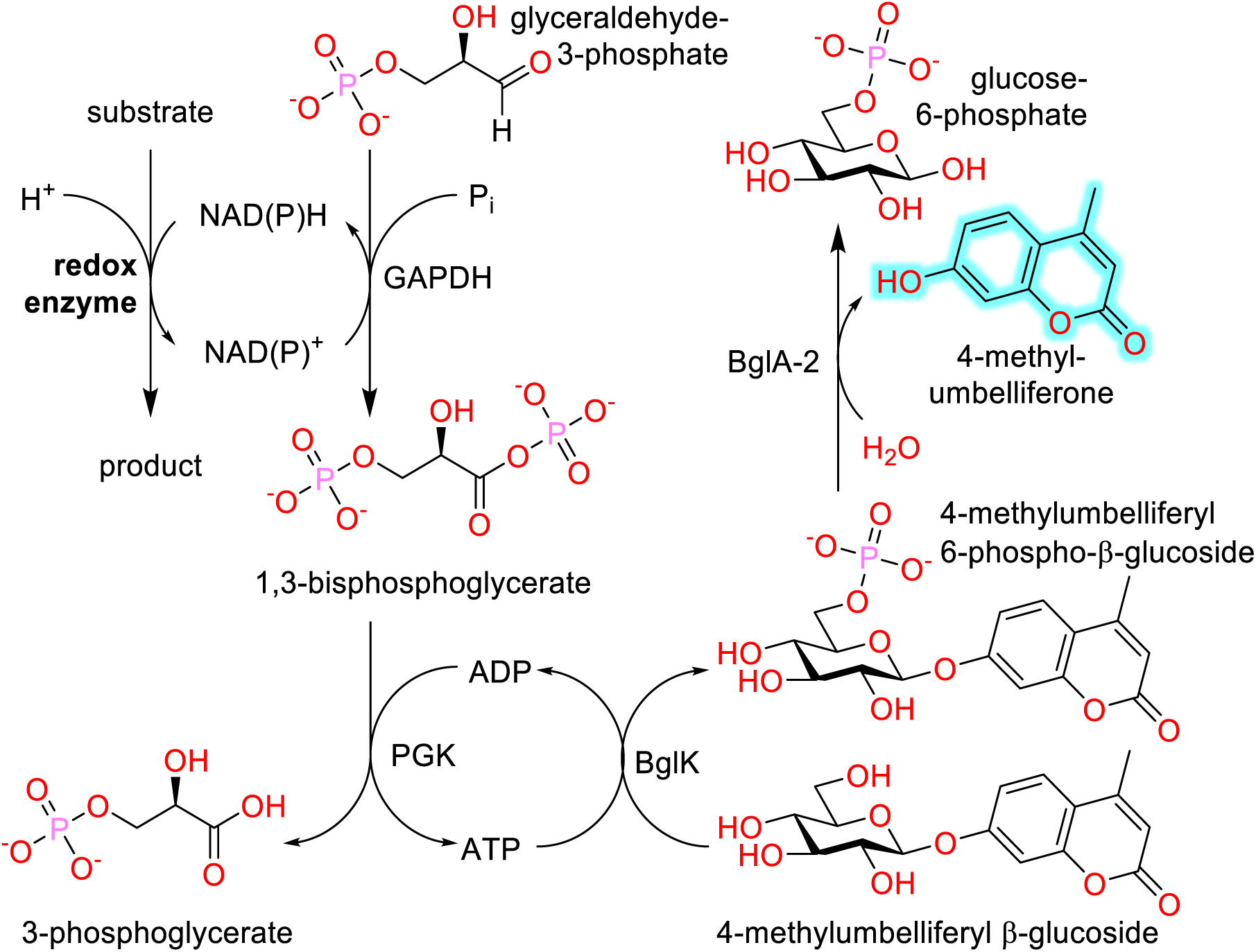
Fluorescence-based, enzyme cascade-coupled assay to screen activity of oxidoreductase (redox) enzymes.

### Validating the downstream coupled enzyme cascade to detect released NAD^+^

Prior to coupling the cascade to any redox reaction, we tested whether the series of GAPDH-, PGK-, BglK-, and BglA-2-catalyzed reactions could be used to detect NAD^+^ specifically and sensitively with the release of fluorescent 4-methylumbelliferone. We observed (**Fig. 2**) that the combined reactions can indeed successfully generate a detectable fluorescence output when NAD^+^ is included with other co-substrates (G3P, P_i_, ADP, and MU-Glc) along with the coupling enzymes (each at 20 µg/mL). Several control conditions were also tested excluding various components of the reaction mixture and, notably, no appreciable fluorescence was observed when NAD^+^ was absent, nor when either ADP or G3P were absent. Similarly, the exclusion of the mixture of coupled enzymes resulted in no detectable fluorescence. A strong fluorescence is only observed when all of the components are present. Together these results indicate that the coupled enzyme cascade functions as designed and is dependent upon NAD^+^ to trigger the release of fluorescence through sequential reactions that also consume G3P along with P_i_ and use ADP to generate ATP, which is used to phosphorylate the fluorogenic substrate before it can be cleaved.

**Figure 2.**
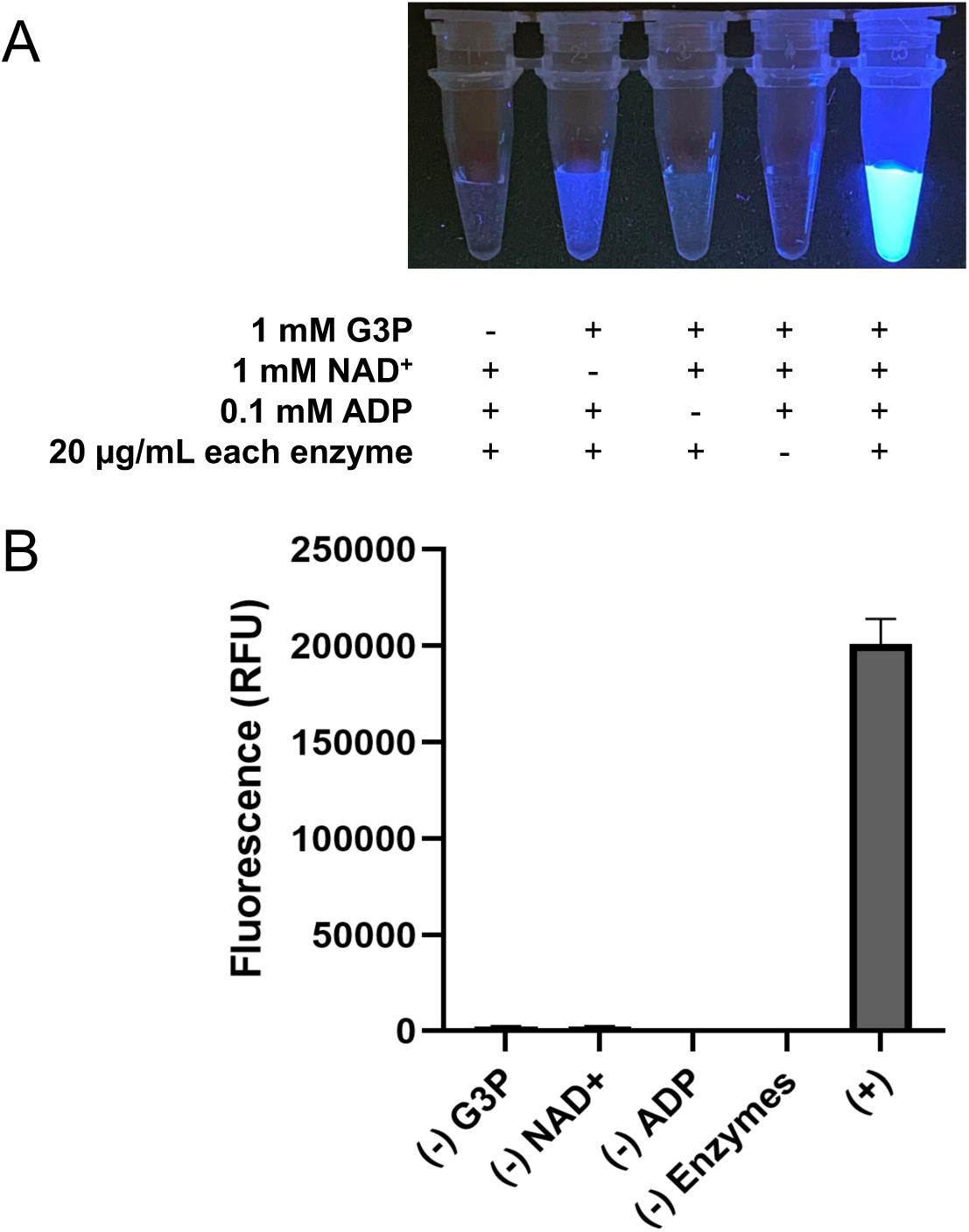
Fluorescence signal resulting from coupled enzyme cascade to detect NAD^+^. (A) Qualitative observation of fluorescence in a reaction including NAD^+^ along with coupled enzymes (GAPDH, PGK, BglK, and BglA-2) and their (co-)substrates (G3P, Pi, ADP, and MU-Glc) but no appreciable fluorescence in controls omitting those components. (B) Quantitative measurement of the fluorescence signal from the coupled enzyme reaction and controls.

### Comparison of absorbance-based assay and fluorescence-based coupled assay for an NADH-dependent redox enzyme (lactate dehydrogenase; LDH)

As a proof of concept, we used our coupled enzyme strategy to test the NADH-dependent reductase activity of lactate dehydrogenase (LDH). NADH-dependent dehydrogenases are conventionally assayed spectrophotometrically. For example, in diagnostics, LDH concentrations in blood samples are measured by monitoring the change of absorbance of NADH at 340 nm [24]. To assay the redox enzyme, we made serial dilutions of LDH, and tested the sensitivity using pyruvate as a substrate. We quantitatively demonstrated the improvement of sensitivity and performance of the coupled enzyme cascade over that of the conventional absorbance-based assay (**Fig 3A, 3B**). The reactions were performed using NADH as the cofactor for LDH-catalyzed reduction of pyruvate to lactate.

**Figure 3.**
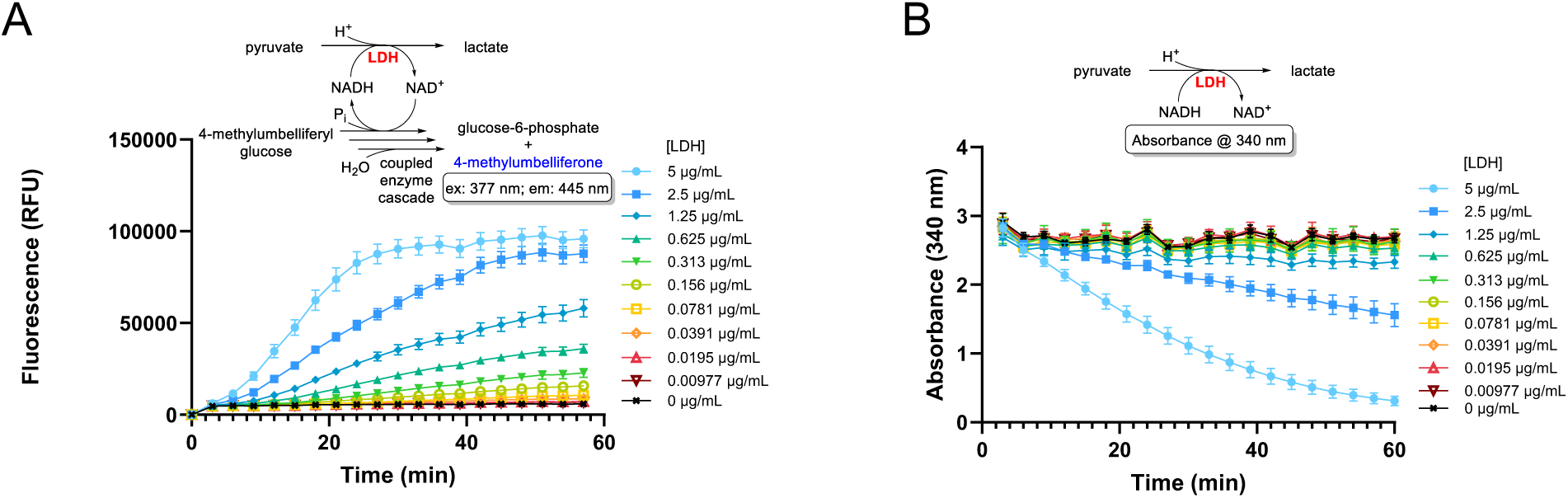
Assay of lactate dehydrogenase. (A) Fluorescence-based assay of the reduction of pyruvate catalyzed by LDH (at varied concentration) with coupled enzyme cascade including coupling enzymes (GAPDH, PGK, BglK, & BglA-2) and their (co)substrates (G3P, NADH, Pi, ADP, MU-Glc). (B) Conventional spectrophotometric assay of the reduction of pyruvate catalyzed by LDH (at varied concentration) by measuring change in NADH absorbance at 340 nm.

The performance was checked by assaying the reaction in multiple replicates over a range of concentrations of LDH and the Z′ (also known as Z-factor, but not to be confused with Z-score) was calculated as a measure of statistical effect size. In high-throughput screening, Z′ is used as a quality indicator for a biochemical assay. Generally, high-throughput screens that exhibit a Z′ between 0.5 and 1 are considered excellent, while a Z′ above 0 but less than 0.5 is considered acceptable. By comparing the continuous fluorescence assay for LDH activity with the classical absorbance-based measurement method, we show that the fluorescence-based assay exhibits greater sensitivity. The Z′ values were determined for the assays with varying LDH concentration, showing that by using the coupled enzyme cascade, the threshold for achieving an excellent assay by Z′ (1 > Z′ ≥ 0.5) could be reached with much lower concentrations of LDH (0.156 µg/mL or 0.66 mU/mL) compared to concentrations required for the conventional absorbance-based assay (5 µg/mL or 21 mU/mL), amounting to a 32-fold difference (**Table 1**). The less stringent threshold of achieving an acceptable assay by Z′ (0.5 > Z′ > 0), could be reached in the coupled enzyme cascade using only 0.039 µg/mL of LDH (0.17 mU/mL) compared with 2.5 µg/mL of LDH (11 mU/mL) needed in the conventional absorbance-based assay (a 64-fold difference).

**Table 1.**
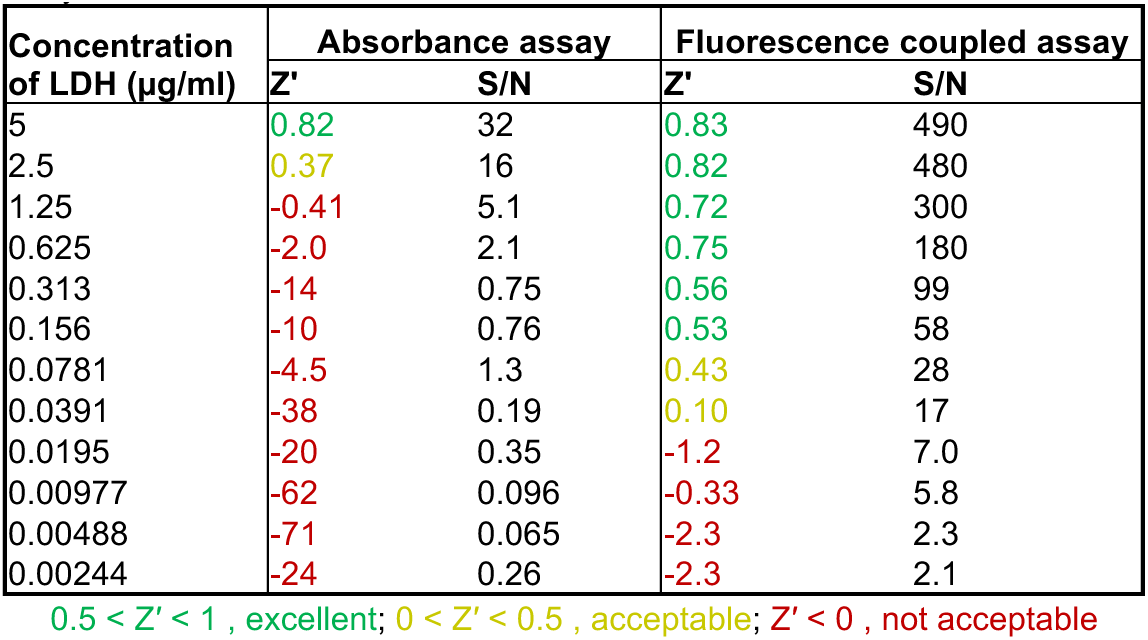
Z’ values and signal-to-noise ratio for LDH assays performed by conventional absorbance assay or fluorescence-based assay by coupled enzyme cascade.

To quantitatively compare the sensitivity of the LDH assay by coupled enzyme cascade to the assay by conventional absorbance measurement, we calculated the signal-to-noise ratio as the mean difference of the test signal and background readings divided by the standard deviation of the background readings. We observed that for a given concentration of LDH, assay by coupled enzyme cascade consistently gave a signal-to-noise ratio that is orders of magnitude greater than assay by conventional absorbance measurements (**Table 1**).

Our strategy also has the advantage that it is effective with a range of NADH cofactor concentrations. Whereas the conventional absorbance-based assay proceeds with a decrease in the concentration of NADH that is measured spectrophotometrically, our fluorescence-based coupled enzyme cascade strategy is designed such that NADH is constantly regenerated by the coupling enzyme, GAPDH (which recycles NAD^+^). Thus, considerably lower concentrations of NADH could be used in LDH assays with the coupled enzyme cascade since the reaction is not limited by the amount of redox cofactor initially supplied. The comparisons of LDH assays by coupled enzyme cascade and by conventional absorbance measurement detailed above were performed with an NADH concentration of 1 mM, which is typical as the upper end of the dynamic range for absorbance measurements at 340 nm (with higher concentrations falling outside the linear range of Beer’s Law). We also tested the assay of LDH by coupled enzyme cascade using considerably lower or considerably higher concentrations of NADH. With 0.1 mM, 1 mM, or 2 mM NADH in the coupled enzyme cascade, the Z′ values remained consistent (**Supplementary Table S1** and **Supplementary Figure S1**), demonstrating that our fluorescence-based assay by coupled enzyme cascade is versatile with a wide range of NADH cofactor concentrations compared to the conventional absorbance-based method.

### A mutant *E. coli* GAPDH enzyme with dual NAD^+^/NADP^+^ specificity is suitable as a coupling enzyme to assay NADPH-dependent oxidoreductases

While we successfully demonstrated the sensitive detection of NADH-dependent oxidoreductase activity using lactate dehydrogenase (LDH), many important redox enzymes, such as Old Yellow Enzymes (OYEs), preferentially use NADPH as a cofactor instead of NADH. In our initial studies, we noted that the GAPDH from *E. coli* (GapA) employed in our coupled enzyme cascade has a strong preference for NAD^+^ over NADP^+^ (**Fig. 4**). We therefore sought to modify our enzyme cascade to allow us to sensitively detect the activity of oxidoreductases that use NADPH instead of NADH. Slivinskaya *et al*. reported a GapA double mutant (G188T+P189K) that showed significantly higher activity on NADP^+^ compared to the wild-type enzyme [25]. We generated the GapA G188T+P189K mutant by site directed mutagenesis and recombinantly expressed and purified the enzyme. We then tested the downstream coupled enzyme cascade (in the absence of oxidoreductase) in detecting NAD^+^ or NADP^+^, either of which was added to start a reaction containing a GAPDH which was either the wild-type or mutant (G188T+P189K) *E. coli* GapA enzyme, together with PGK, BglK, and BglA-2 and with the necessary substrates for these coupling enzymes (G3P, P_i_, ADP, MU-Glc). The reactions were allowed to continue for 10 minutes before termination, after which fluorescence was measured at 445 nm. For the GapA mutant, **Fig. 4** shows the stark effect of the G188T+P189K double substitution on the fluorescence observed when NADP^+^ is supplied to trigger the downstream coupled enzyme cascade. While activity on NAD^+^ remains similar, the ability of GapA G188T+P189K to utilize NADP^+^ is markedly increased compared to the wild-type enzyme, leading to a much higher release of fluorescence in the enzyme cascade. Although the GapA mutant still exhibits a preference for NAD^+^, its relaxed specificity allows it to use NADP^+^ to trigger the coupled enzyme cascade, which enables coupled assay of NADPH-dependent redox enzymes with sensitive release of fluorescence.

**Figure 4.**
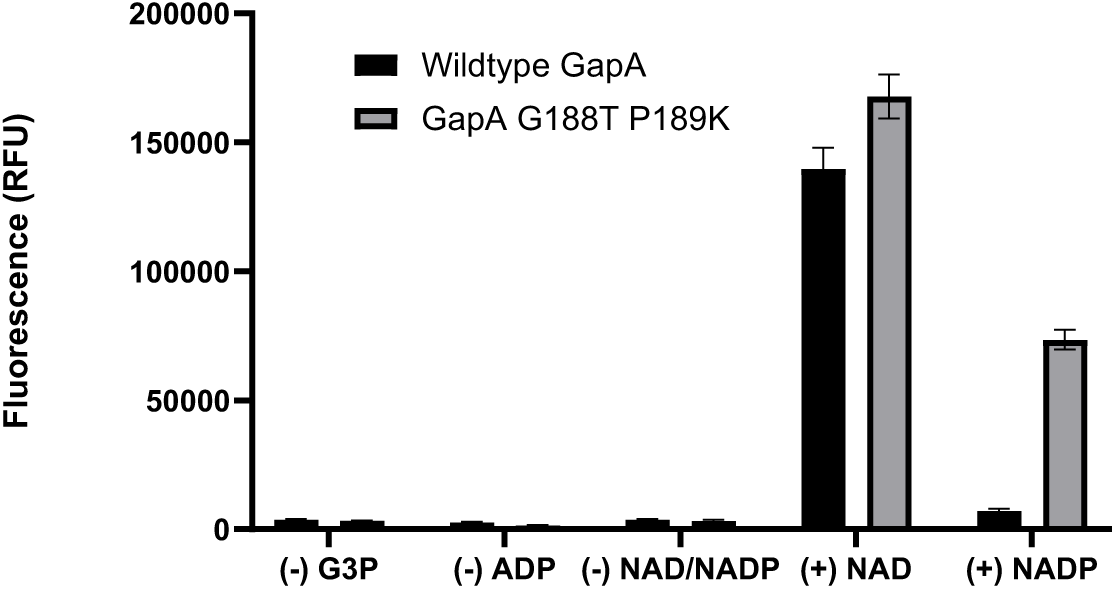
Fluorescence signal of the coupled enzyme cascade using either the wild type GAPDH from *E. coli* (GapA), or the mutant version (GapA G188T+P189). Full reactions include GAPDH (wild-type or mutant), PGK, BglK, and BglA-2, together with substrates G3P, ADP, and either NAD^+^ or NADP^+^, while negative (-) control reactions lack G3P, ADP, or NAD(P)^+^.

### Comparison of absorbance-based assay and fluorescence-based coupled assay for an NADPH-dependent redox enzyme (Old Yellow Enzyme II; OYE2)

Having modified the coupled enzyme cascade with a variant GAPDH (mutant *E. coli* GapA G188T+P189K) so that the sequential reactions can be triggered by NADP^+^, we then aimed to use our strategy to assay the activity of an NADPH-dependent oxidoreductase. We were interested in testing the activity of Old Yellow Enzymes (OYEs) because of their relevance in green chemistry and biocatalysis. Many OYEs tend to prefer NADPH as a cofactor, which they use as a donor of electrons that are transferred *via* a tightly bound flavin mononucleotide (FMN) electron carrier to reduce C=C double bonds of the enzyme’s substrate (**Fig. 5A**) [22, 26, 27]. Using our modified cascade with the engineered GAPDH, we coupled the sequential reactions to OYE2 activity to perform a fluorescence-based assay, which we compared to the conventional absorbance-based assay of OYE2. The setup to test either assay strategy with OYE2 was similar to what we had done with LDH, with a key distinction being that NADPH was used as the redox cofactor to reduce a 2-cyclohexenone substrate in assays of OYE2 activity, which—in the fluorescence-based assay— could be detected by the use of *E. coli* GapA G188T+P189K as the GAPDH enzyme in the coupled enzyme cascade (that also included PGK, BglK and BglA-2). In our initial tests of NADPH-dependent OYE2 activity, we indeed observed that the performance and sensitivity (determined by quantifying Z′ and S/N for different concentrations of OYE2 tested) was better in the fluorescence-based assay by coupled enzyme cascade compared to the conventional assay by absorbance, but the difference was not as stark as we had observed for our assays with LDH (**Supplementary Figure S2**, and **Supplementary Table S2**). Examination of the fluorescence curves in those initial tests revealed the likely reason for this; it appeared that for higher concentrations of OYE2 tested, the OYE2-catalyzed reaction was no longer the rate-limiting-step in the enzyme cascade which included 20 µg/mL of each coupling enzyme. This was consistent with our observation that the GAPDH enzyme in these assays, *E. coli* GapA G188T+P189K, has lower activity on NADP^+^ than it (or the wild-type GapA) does on NAD^+^ leading to a lower fluorescence signal in the coupled enzyme cascade (**Fig. 4**). To address this, we further modified the coupled enzyme cascade, increasing the concentration of GapA G188T+P189K ten-fold to 200 µg/mL (keeping PGK, BglK and BglA-2 at 20 µg/mL) and we observed that this alleviated the bottleneck at the GAPDH-catalyzed reaction, resulting in increased signal and sensitivity by fluorescence (**Fig. 5B**). This accounted for an overall improvement in performance as quantified by Z′ (**Table 2**).

**Figure 5.**
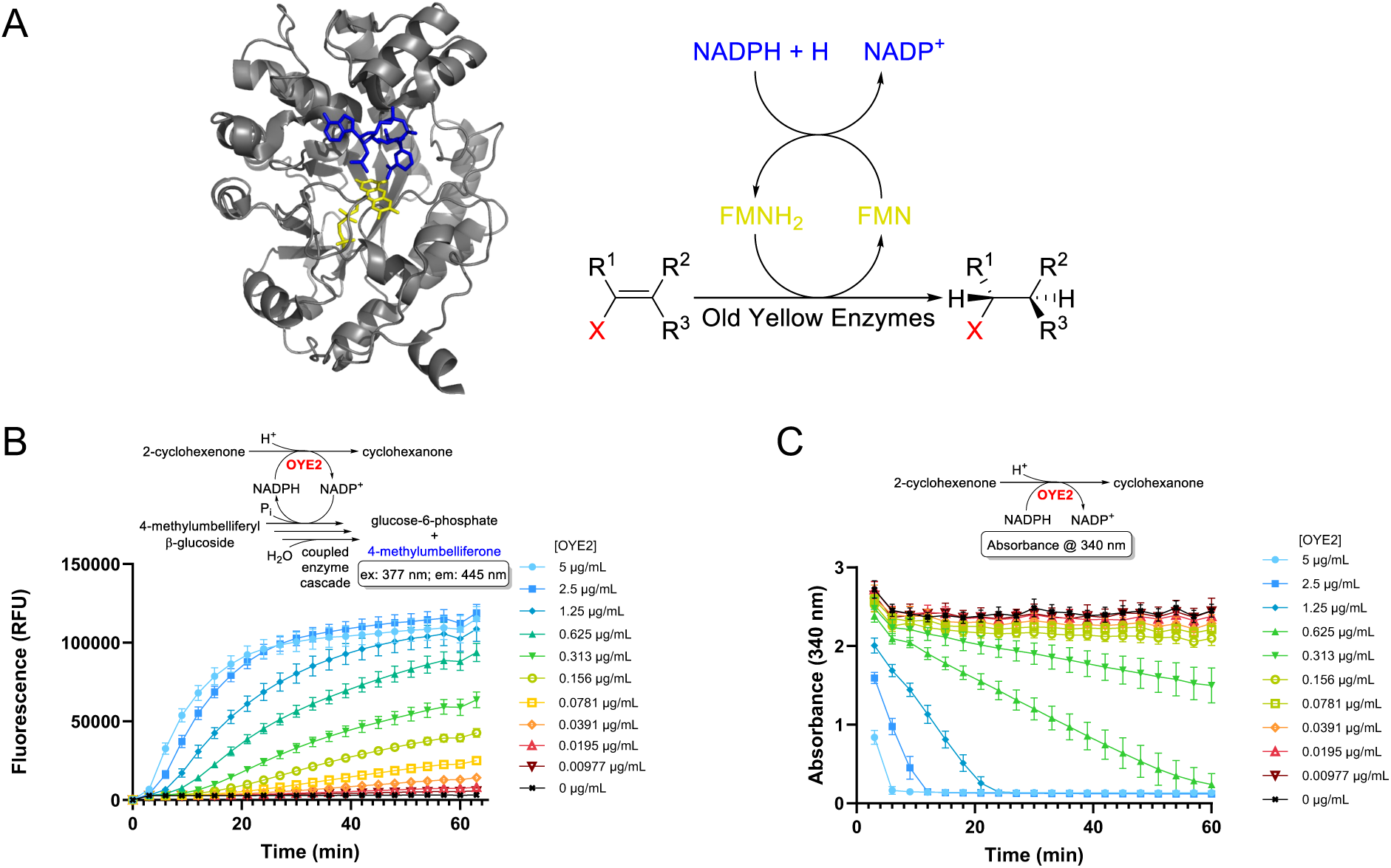
Assay of Old Yellow Enzyme-catalyzed reactions. (A) Left: model of the ternary structure of the OYE2 enzyme in complex with FMN and NADPH cofactors generated by docking NADPH to the crystal structure of the OYE2 and FMN complex (PDB: 9FH7); right: reaction scheme illustrating the reduction of a C=C double bond (ene functionality) catalyzed by Old Yellow Enzymes. (B) Fluorescence-based assay of the reduction of 2-cyclohexenone catalyzed by OYE2 (at varied concentration) with coupled enzyme cascade including coupling enzymes (GapA G188T+P189K, PGK, BglK, & BglA-2) and their (co)substrates (G3P, NADPH, P_i_, ADP, MU-Glc). (C) Conventional spectrophotometric assay of the reduction of 2-cyclohexenone catalyzed by OYE2 (at varied concentration) by measuring change in NADPH absorbance at 340 nm.

**Table 2.** Z’ values and signal-to-noise ratio for OYE2 assays performed by conventional absorbance assay or fluorescence-based assay by coupled enzyme cascade.

| Concentration of OYE2 ($\mu\text{g/mL}$) | Absorbance assay | | Fluorescence coupled assay | |
| --- | --- | --- | --- | --- |
| | $Z'$ | S/N | $Z'$ | S/N |
| 5 | 0.87 | 27 | 0.79 | 670 |
| 2.5 | 0.89 | 28 | 0.85 | 690 |
| 1.25 | 0.89 | 28 | 0.77 | 630 |
| 0.625 | 0.70 | 27 | 0.80 | 540 |
| 0.313 | 0.036 | 11 | 0.74 | 360 |
| 0.156 | -0.48 | 4.2 | 0.77 | 230 |
| 0.0781 | -0.98 | 2.9 | 0.68 | 130 |
| 0.0391 | -2.6 | 1.9 | 0.67 | 65 |
| 0.0195 | -6.9 | 0.72 | 0.54 | 29 |
| 0.00977 | -370 | 0.024 | 0.50 | 15 |
| 0.00488 | -4.9 | 1.4 | 0.13 | 8.7 |
| 0.00244 | -5.1 | 0.89 | -0.088 | 6.5 |

0.5 < $Z'$ < 1 , excellent; 0 < $Z'$ < 0.5 , acceptable; $Z' < 0$ , not acceptable

For OYE2 activity, compared to the conventional assay by absorbance (**Fig. 5C** and **Table 2**), the optimized fluorescence assay by coupled enzyme cascade was far superior in sensitivity and performance. The threshold for achieving an excellent assay by Z′ (1 > Z′ ≥ 0.5) could be reached with a minimum concentration of 9.77 × 10^-3^ µg/mL of OYE2 (0.22 mU/mL) in the optimized assay by coupled enzyme cascade, compared to a minimum of 0.625 µg/mL of OYE2 (14.3 mU/mL) in the conventional assay by absorbance (amounting to a 64-fold difference). The less stringent threshold of achieving an acceptable assay by Z′ (0.5 > Z′ > 0) could be reached with a minimum concentration of 4.88 × 10^-3^ µg/mL of OYE2 (0.11 mU/mL) in the optimized assay by coupled enzyme cascade, compared to a minimum of 0.3125 µg/mL of OYE2 (7.15 mU/mL) in the conventional assay by absorbance (again a 64-fold difference). The signal-to-noise ratio in the assay of OYE2 by the optimized coupled enzyme cascade was also observed to generally be orders of magnitude higher than for the conventional assay by absorbance (**Table 2**).

Together, these results demonstrate that through the use of an enzyme cascade that includes an NAD^+^/NADP^+^ dual-specific GAPDH (along with PGK, BglK, and BglA-2), our strategy allows for versatility to detect NADPH-dependent as well as NADH-dependent oxidoreductase activity in a manner that is superior to the conventional absorbance-based assay.

### Adaptations to economize the assay by coupled enzyme cascade: *in situ* G3P generation and use of a nanodroplet-compatible fluorogenic substrate

Although we demonstrated that we can assay both NADH- and NADPH-dependent oxidoreductases with high performance and high sensitivity using our coupled enzyme cascade, one major concern in employing any assay strategy for high-throughput screening is cost. The ability to detect activity with a very small amount of enzyme through a strong fluorescence signal is certainly favourable for an economical screen accounting for an estimated cost of CAD $0.55 per assay using our coupled enzyme cascade strategy. The largest contributor to this cost is G3P, a co-substrate for GAPDH which acts as the first coupled enzyme in the cascade. In an effort to reduce the cost of the assay, we tested whether we could produce this expensive reagent *in situ* from an inexpensive precursor. We aimed to extend the coupled enzyme cascade so that G3P could be produced from cheaply available fructose-1,6-bisphosphate (FBP) through fructose bisphosphate aldolase (FBA)- and triose phosphate isomerase (TPI)-catalyzed reactions. First, we tested whether an extended enzyme cascade including FBA and TPI (in addition to the typical GAPDH, PGK, BglK, and BglA-2 at 20 µg/mL each) could detect NAD^+^ with the release of fluorescent 4-methylumbelliferone in a reaction that also contained (co-)substrates FBP, P_i_, ADP, and MU-Glc (**Supplementary Figure S3**). Indeed, fluorescence was observed in the reaction mixture in which FBP, but no exogenous G3P was added (while no appreciable fluorescence was observed in controls lacking FBP, NAD^+^, P_i_, ADP, or the enzyme mix), indicating that *in situ* generation of G3P from FBA- and TPI-catalyzed conversion of FBP completes the reaction cascade. We then tested the assay of LDH activity by coupling it to the extended cascade (including FBA and TPI) and observed high sensitivity and performance comparable to the LDH assay by our standard coupled enzyme cascade (**Supplementary Figure S3** and **Supplementary Table S3**). Bypassing the need to commercially source G3P as a starting co-substrate (and generating it *in situ* from cheaply available F6P instead) significantly decreases the cost of the assay, reducing it from CAD $0.55 per assay to CAD $0.17 per assay (**Supplementary Table S4**).

Another important step to economizing high-throughput screens is miniaturization. While we have optimized the performance of our assay by coupled enzyme cascade on the microliter-scale in 384-well microplates, further scaling down assay volumes to the nanoliter-scale would increase savings dramatically. Droplet-based techniques using microfluidics have emerged as ultra high throughput strategies for screening mutant libraries in directed evolution. To screen enzyme activity in such an experimental setup, the fluorescence read-out should be detected from assays carried out in nanodroplets in either a water-in-oil or a water-in-oil-in-water emulsion with minimal signal loss or cross-talk caused by the fluorophore leaking out of the aqueous droplets. Despite being a useful fluorescent reporter for microplate-based screening, 4-methylumbelliferone can diffuse between aqueous droplets and the oil phase of an emulsion due to its hydrophobicity, resulting in poor read-outs and bleed through of the signal in droplet-based systems. To address this, Withers and colleagues had previously synthesized a hydroxycoumarin-based fluorophore called Jericho Blue, specialized for emulsion droplet-based screening of glycosidases [28]. Thus, to explore the potential of adapting our coupled enzyme cascade strategy to an emulsion droplet-based system, we tested the use of Jericho Blue as a reporter molecule instead of 4-methylumbelliferone. Using a β-glucoside of Jericho Blue (JB-Glc) to replace MU-Glc in our standard conditions for LDH assays by coupled enzyme cascade, we observed that we could detect activity by Jericho Blue fluorescence (390 nm excitation, 450 nm emission), indicating that JB-Glc can take part in the cascade where BglK can catalyze its ATP-dependent conversion to Jericho Blue 6-phospho-β-glucoside, which can then be cleaved by BglA-2-catalyzed hydrolysis to release the fluorescent Jericho Blue reporter molecule (**Supplementary Figure S4**). We also observed that assay characteristics, Z′ and S/N, were very good using the coupled enzyme cascade with JB-Glc and similar to those for the coupled enzyme cascade using MU-Glc (**Supplementary Table S5**).

## Conclusions and Future Perspectives

Developing a high-throughput screen is a key step in enzyme engineering. In directed evolution, screening large libraries of enzyme variants demands assays that are both convenient and highly sensitive to detect even small improvements in activity. Microplate-based assays are typically used for screening thousands of variants, while droplet-based microfluidic systems are necessary for libraries with millions of variants. In both formats, a sensitive and reliable readout is critical for efficiently identifying rare, high-performing enzymes and accelerating the optimization process. The fluorescence-based coupled enzyme assay developed in this study is an improved high-throughput method that can be employed toward engineering NAD(P)H/NAD(P)^+^-dependent oxidoreductases. The assay platform is built on a cascade of enzymes that ultimately generates a strong fluorescence signal upon the hydrolysis of 4-methylumbelliferyl-β-D-6-phosphoglucoside. By measuring fluorescence generated by the enzyme cascade instead of relying on the conventional absorbance of NAD(P)H, our assay strategy achieves superior performance and sensitivity, as demonstrated by statistical measures of assay quality such as the Z′ factor and signal-to-noise ratio, particularly at low enzyme concentrations.

Our strategy was initially effective for assaying NADH-dependent oxidoreductases, as demonstrated with LDH. A major advancement of this study was adapting the coupled enzyme cascade to also enable detection of NADPH-dependent enzymes, which we validated using OYE2—a member of the Old Yellow Enzyme family with important industrial and green chemistry applications. By incorporating a dual-specific, engineered GAPDH capable of recognizing both NAD^+^ and NADP^+^, we expanded the scope of our system to allow sensitive screening of both NADH- and NADPH-dependent redox enzymes.

To further improve the economy and scalability of our coupled enzyme cascade assay for high-throughput screening, we developed strategies to reduce reagent costs and adapt the assay for miniaturized formats. By extending the cascade to generate G3P in situ from inexpensive FBP using FBA and TPI, we significantly lowered the assay cost from CAD $0.55 to $0.17 per reaction, offering substantial savings when screening large libraries. To enable future adaptation to droplet-based microfluidic screening, we replaced 4-methylumbelliferone with the more hydrophilic Jericho Blue fluorophore, maintaining strong assay performance while minimizing potential signal leakage in emulsion systems. Together, these advances make our fluorescence-based coupled enzyme cascade a versatile, cost-effective, and scalable platform for the directed evolution of oxidoreductases.

In conclusion, this study presents a versatile and sensitive fluorescence-based coupled-enzyme assay for detecting NAD(P)H/NAD(P)^+^-dependent redox enzyme activity. Enhancements to improve the assay’s sensitivity, cost-efficiency, and adaptability to droplet-based systems lay the foundation for its broader application in enzyme discovery and optimization.

## Materials and Methods

### Cloning and construction of plasmids

Details of the plasmid constructs are summarized in **Supplementary Table S6**.

Expression plasmids (pTrcHis-BglK and pET28-SpBglA-2) coding the β-glucoside kinase (BglK) from *Klebsiella pneumoniae* and the 6-phospho-β-glucosidase (BglA-2) from *Streptococcus pnemoniae* TIGR4 were previously constructed as described in literature [29, 30]. The plasmids carrying the genes *gapA*, *pgk*, *fbaA*, and *tpiA* from *E. coli*, coding for the glyceraldehyde 3-phosphate dehydrogenase (GAPDH), phosphoglycerate kinase (PGK), fructose-1,6-bisphosphate aldolase (FBA), and triose phosphate isomerase (TPI) enzymes, respectively, and the plasmid carrying the *OYE2* gene from *S. cerevisiae*, coding for Old Yellow Enzyme II, were each constructed by cloning the target gene in a pET-28a expression vector as follows. Genomic DNA from *E. coli* K12 or *S. cerevisiae* was used as the template for PCR reactions in which the desired genes were amplified with flanking regions of homology pET-28a vector, which were included in the 5′-ends of the forward and reverse primers (listed in **Supplementary Table S7**). To insert each of the PCR-amplified genes into the vector, pET-28a was first linearized by digestion with BamHI and NdeI and purified using an EZ-10 Spin Column PCR Products Purification Kit (BioBasic), then this was combined with PCR-amplified DNA containing the target gene flanked with sequences homologous to the ends of the linearized vector and assembled by the “Seamless Ligation Cloning Extract” method (SLiCE) [31]. The resulting plasmids, pET28-gapA, pET28-pgk, pET26-fbaA, pET28-tpiA and pET28-OYE2 would later be used for recombinant expression of proteins with *N*-terminal hexahistidine (His_6_) tags.

The plasmid coding the GapA G188T+P189K mutant version of the *E. coli* GAPDH enzyme was generated by PCR-based site-directed mutagenesis using pET28-gapA as a template and the mutagenic primers indicated in **Supplementary Table S7**.

### Preparation of proteins

Lactate dehydrogenase (LDH) was commercially sourced. All other proteins described were recombinantly expressed in *E. coli*. The protocols for expression and purification of BglK and BglA-2 are reported elsewhere [29, 30], but for the remaining recombinantly expressed enzymes, we performed the following general procedure. Expression plasmids were used to transform *E. coli* BL21(DE3). From each transformation, a single colony was used to inoculate a seed culture, grown at 37 °C overnight in 10 mL of Terrific Broth (TB), that was then used to inoculate an expression culture in 800 mL of TB media supplemented with the appropriate antibiotic (50 µg/mL kanamycin for plasmids derived from pET-28a). The expression culture was then grown at 37 °C in a shaking incubator at 220 rpm until it was induced at log phase (OD_600_ = 0.4∼0.8) with the addition of IPTG to a final concentration of 0.5 mM. The induced culture was then grown overnight, at 18 °C in a shaking incubator.

For purification of each of the His_6_-tagged proteins, the cultures grown overnight were first harvested by centrifugation at 10,000 *g* for 15 minutes at 4 °C, then the pellet was resuspended in 30 mL of wash/resuspension buffer (50 mM Tris, 500 mM NaCl, 10 mM imidazole, pH 7.5), supplemented with DNAse I, RNAse A and lysozyme added to a final concentration of 5 µg/mL and the addition of one EDTA-free protease inhibitor tablet (Roche) per 30 mL. The cells were lysed by sonication (25% amplitude, pulse on 5 for seconds, pulse off 15 for seconds, 3 minutes total pulse time) and centrifuged to remove cell debris. Insoluble material was removed from the crude lysate by centrifugation at 15,000 *g* for 30 minutes at 4 °C and the supernatant was filter-sterilized using a 0.22-µm filter and applied to a nickel nitrilotriacetate (Ni-NTA) agarose resin column (1 mL) pre-equilibrated with wash buffer. Protein fractions were eluted by AKTA-FPLC with an increasing gradient of imidazole (10 mM to 300 mM). The fractions were analysed on SDS-PAGE and those containing the desired protein were pooled and concentrated to 3 mL using a 30,000 Da molecular weight cut-off Vivaspin 6 or Vivaspin 20 concentrator (Sartorius). Buffer exchange was performed using a 10DG desalting column (Bio-Rad) that was pre-equilibrated with storage buffer (50 mM Tris, 150 mM NaCl, pH 7.5), following the manufacturer’s guidelines. The final concentration of purified protein was determined by Bradford assay or by bicinchoninic acid (BCA) assay. Small aliquots of purified protein were flash frozen using liquid nitrogen and stored at −80 °C.

### Test of the coupled enzyme cascade to detect NAD^+^ or NADP^+^

Assays were set up in which the complete reaction was carried out in solution of phosphate-buffered saline (pH 7.2) containing 1 mM MgCl_2_, 1 mM MU-Glc, 1 mM G3P, 0.1 mM ADP, and 1 mM of either NAD^+^ or NADP^+^ and reactions were started with the addition of an enzyme mix consisting of a GAPDH (either wild-type *E. coli* GapA or the G188T+P189K mutant), PGK, BglK, and BglA-2 each to a final concentration of 20 µg/mL. Control reactions were also carried out in which G3P, ADP, NAD(P)^+^, or the enzyme mixture were omitted. Reactions in 10 µL volumes were incubated for 5 to 10 minutes at 25 °C then terminated with the addition of 50 µL stop buffer (0.1 M glycine, pH 10.3) before measuring fluorescence (excitation at 365 nm, emission at 445 nm).

An alternative version of the coupled enzyme cascade was tested in which G3P was omitted and instead 1 mM F6P was included, with the addition of 20 µg/mL of each of FBA and TPI to the enzyme mixture (facilitating the *in situ* generation of G3P from F6P).

### Continuous fluorescence-based assays of NAD(P)-dependent oxidoreductases by coupled enzyme cascade

The fluorescence of 4-methylumbelliferone in a 50 µL reaction was measured in a 384-well plate using Clariostar plate reader (BMG) with automated dispensing. Typically, the reaction was carried out in a solution containing phosphate-buffered saline (pH 7.2) and 1 mM MgCl_2_, 1 mM MU-Glc, 1 mM G3P, 0.1 mM ADP and a coupling enzyme mix containing 20 µg/mL each of GAPDH (either wild-type *E. coli* GapA or the G188T+P189K mutant), PGK, BglK, and BglA-2. Some experiments were performed with GapA G188T+P189K at 200 µg/mL. These components were added with redox enzyme (either LDH or OYE2) along with its substrate (either pyruvate or 2-cyclohexenone at 1 mM). Prior to the start of the reaction, the mixture was incubated at 37 °C in the absence of NAD(P)H for at least 5 minutes. The reaction was then started by adding either NADH or NADPH to a final concentration of 1 mM. The fluorescence was measured over 1 hour (time points every 3 minutes) with excitation at 372 nm and emission at 445 nm.

An alternative version of the continuous fluorescence assay by coupled enzyme cascade was tested in which G3P was omitted and instead 1 mM F6P was included, with the addition of 20 µg/mL of each of FBA and TPI to the enzyme mixture (facilitating the *in situ* generation of G3P from F6P).

Yet another alternative version of continuous fluorescence assay by coupled enzyme cascade was tested in which JB-Glc (provided as a generous gift by S. G. Withers) was used as the fluorogenic substrate instead of MU-Glc. The fluoresecence of Jericho Blue was monitored at 390 nm excitation, 450 nm emission.

A ClarioStar plate reader (BMG) with automated dispensing was used to perform the experiment. For the calculation of Z′ for assays, eight replicates were performed per enzyme concentration and control condition in an experiment performed on a single 384-well microtitre plate.

### Continuous enzyme assays of NAD(P)H-dependent oxidoreductases by monitoring NAD(P)H absorbance at 340 nm

The absorbance of a 50 µL reaction was measured in a 384-well plate using Clariostar plate reader (BMG) with automated dispensing. The reaction was carried out at (pH = 7.2), with final reaction components at 1 mM MgCl_2,_ 1 mM substrate (pyruvate for LDH or 2-cyclohexone for OYE2) and varying concentrations of redox enzyme (LDH or OYE2). Each concentration was tested in 8 replicates. The redox enzyme and substrate were pre-incubated at 37 °C for at least 5 minutes. The reaction was started by adding 1 mM NAD(P)H final concentration (NADH was added for LDH and NADPH was added for OYE2). The absorbance was measured at 340 nm over 1 hour (time points every 3 minutes).

For the calculation of Z′ for assays, eight replicates were performed per enzyme concentration and control condition in an experiment performed on a single 384-well microtitre plate.

### Calculation of assay parameters

Z′ was calculated using the following formula where *σ*^+^ is the standard deviation of the test signal, *σ*^-^ is the standard deviation of the background readings (negative control), µ^+^ is the mean of the test signal, and µ^-^ is the mean of the background readings (negative control).

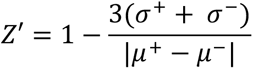

For both the conventional absorbance-based assays and the fluorescence-based assay by coupled enzyme cascade, we observed that Z′ calculated from the endpoint values of the last reading (absorbance or fluorescence) of the 1-hour assay were consistently higher than those calculated from the slopes of the absorbance or fluorescence curves, and thus we recorded Z′ calculated from endpoint values in our results.

The signal-to-noise ratio (S/N) was calculated by the following formula, where µ^+^, µ^-^, and σ^-^ are defined as above.

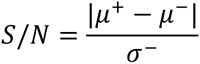

### Cost analysis of assay by coupled enzyme cascade

Pricing from specific vendors used in the cost analysis is summarized in **Supplementary Table S4**. The cost of recombinant enzymes produced in-house was estimated to be CAD $25 per mg based on the time and material for lab personnel to express and purify proteins in batches on the scale of 10 mg.

## Contributions

The work was conceived by DHK. JS and DHK cloned the genes of enzymes and constructed expression vectors. TG, JS, and DHK recombinantly expressed and purified proteins. TG and DHK performed endpoint assays to validate detection of NAD^+^ by coupled enzyme cascade. JS further developed and optimized the continuous assay by coupled enzyme cascade. TG performed experiments with continuous assays of LDH and OYE2 enzymes to determine assay parameters and statistics. TG performed experiments with adaptations meant to economize the fluorescence assay by continuous coupled enzyme cascade. The manuscript was written by TG and DHK.

## Acknowledgments

This work was financially supported by the Natural Sciences and Engineering Research Council of Canada through a Discovery Grant to DHK (grant no. RGPIN-2023-04793) and through an Alexander Graham Bell Canada Graduate Scholarship to JS (application ID 575702-2022). We thank S. G. Withers for the kind gift of Jericho Blue β-glucoside (JB-Glc) used in our experiments.

