## Supplementary for "A high sensitivity strategy to screen NAD(P)H-dependent oxidoreductase activity by coupled enzyme cascade"

**Supplementary Information for "A high sensitivity strategy to screen NAD(P)H-dependent oxidoreductase activity by coupled enzyme cascade"**

Trisha Ghosh[1,†], Jacob Sicheri[1,†], David H. Kwan[1,2,*]

[1]Department of Biology and [2]Department of Chemistry and Biochemistry, Concordia University, 7141 Sherbrooke Street West, Montreal, Quebec, Canada, H4B 1R6

† These authors contributed equally

**Supplementary Table S1**. Z’ values and signal-to-noise ratio for LDH assays performed by conventional absorbance assay or fluorescence-based assay by coupled enzyme cascade with different concentrations of NADH

| | Absorbance | | Fluorescence | | | | | |
|---|---|---|---|---|---|---|---|---|
| | 1 mM NADH | | | | 0.1 mM NADH | | 2 mM NADH | |
| **Concentration of LDH (µg/ml)** | **Z'** | **S/N** | **Z'** | **S/N** | **Z'** | **S/N** | **Z'** | **S/N** |
| 5 | 0.82 | 32 | 0.83 | 490 | 0.80 | 130 | 0.33 | 120 |
| 2.5 | 0.34 | 16 | 0.82 | 480 | 0.88 | 150 | 0.71 | 130 |
| 1.25 | -0.41 | 5.1 | 0.72 | 300 | 0.41 | 140 | 0.77 | 70 |
| 0.625 | -2.0 | 2.1 | 0.75 | 180 | 0.86 | 150 | 0.72 | 51 |
| 0.313 | -14 | 0.75 | 0.56 | 99 | 0.74 | 100 | 0.51 | 29 |
| 0.156 | -10 | 0.76 | 0.53 | 56 | 0.50 | 66 | 0.15 | 18 |
| 0.0781 | -4.5 | 1.3 | 0.43 | 28 | 0.60 | 38 | 0.25 | 8.5 |
| 0.0391 | -38 | 0.19 | 0.10 | 17 | 0.23 | 22 | 0.036 | 5.0 |
| 0.0195 | -20 | 0.35 | -1.19 | 7.0 | 0.39 | 9.6 | -2.9 | 1.8 |
| 0.00977 | -62 | 0.096 | -0.33 | 5.8 | -0.43 | 7.5 | -0.96 | 2.0 |
| 0.00488 | -71 | 0.065 | -2.26 | 2.3 | -2.3 | 3.0 | -1.7 | 1.4 |
| 0.00244 | -24 | 0.26 | -2.34 | 2.1 | -2.6 | 1.8 | -1.6 | 1.4 |

Z' = 1, ideal ; 0.5 < Z' < 1 , excellent ; 0 < Z' < 0.5 , acceptable ; Z' < 0 , not acceptable

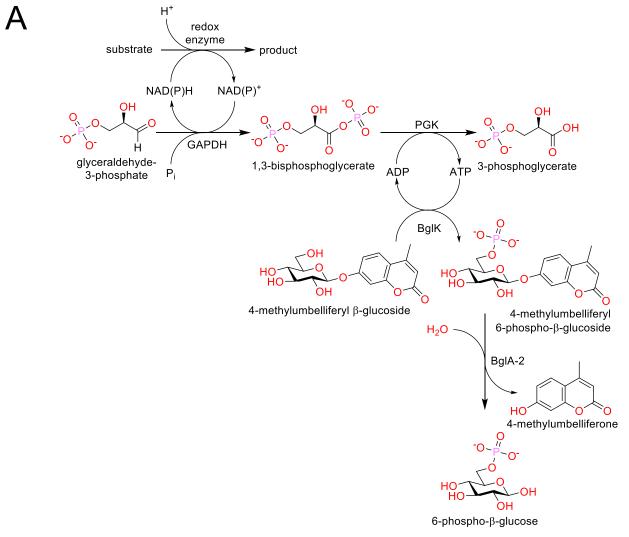


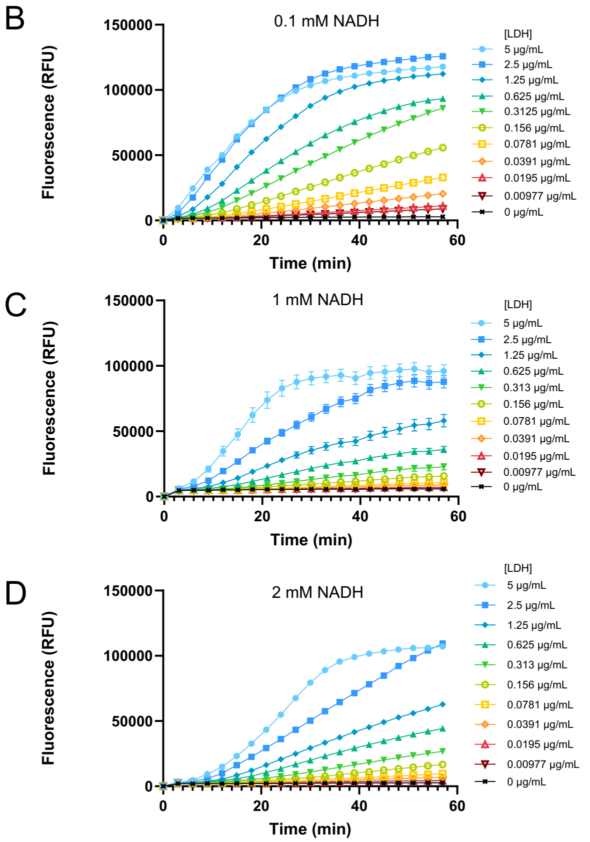


**Supplementary Figure S1.** Scheme of the standard coupled enzyme cascade (A), with which fluorescence-based assays of the reduction pyruvate catalyzed by LDH (at varied concentration) were performed using NADH included at different concentrations: 0.1 mM NADH (B), 1 mM NADH (C), or 2 mM NADH (D).

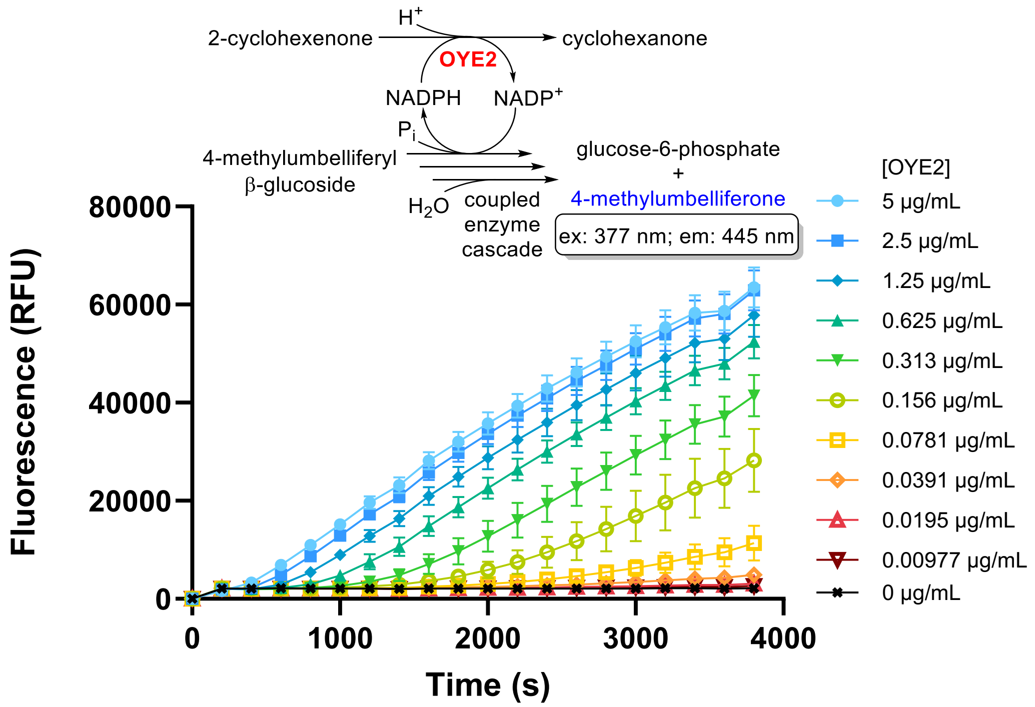


**Supplementary Figure S2.** Fluorescence-based assay of the reduction of 2-cyclohexenone catalyzed by OYE2 (at varied concentration) with coupled enzyme cascade including coupling enzymes (GapA G188T+P189K, PGK, BglK, & BglA-2), at 20 μg/mL each, along with their (co)substrates (G3P, NADPH, $P_i$, ADP, MU-Glc).

**Supplementary Table S2**. Z' values and signal-to-noise ratio for OYE2 assays performed by fluorescence-based assay using a coupled enzyme cascade (including 20 μg/mL each of GapA G188T+P189K, PGK, BglK, and BglA-2)

| **Concentration of OYE2 (μg/ml)** | **Z'** | **S/N** |
|---|---|---|
| 5 | 0.75 | 390 |
| 2.5 | 0.82 | 380 |
| 1.25 | 0.71 | 330 |
| 0.625 | 0.77 | 280 |
| 0.313 | 0.57 | 150 |
| 0.156 | 0.41 | 47 |
| 0.0781 | 0.13 | 12 |
| 0.0391 | -0.35 | 5.0 |
| 0.0195 | -1.1 | 2.8 |
| 0.00977 | -1.4 | 2.3 |
| 0.00488 | -2.7 | 1.6 |
| 0.00244 | -2.3 | 1.8 |

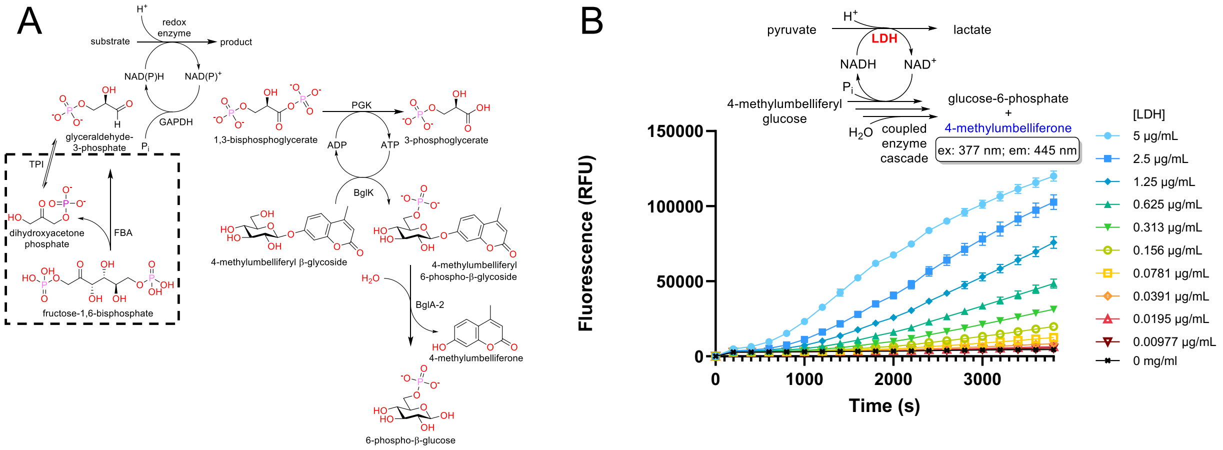


**Supplementary Figure S3.** (A) extended coupled enzyme cascade including FBA and TPI for *in situ* generation of G3P from FBP. Changes from standard coupled enzyme cascade are indicated by dashed line box. (B) Fluorescence-based assay of the reduction pyruvate catalyzed by LDH (at varied concentration) with an extended coupled enzyme cascade including coupling enzymes (FBA, TPI, GAPDH, PGK, BglK, & BglA-2) and their (co)substrates (FBP, NADH, Pi, ADP, MU-Glc).

**Supplementary Table S3**. Z’ values and signal-to-noise ratio for LDH assays performed by fluorescence-based assay using an extended coupled enzyme cascade including coupling enzymes (FBA, TPI, GAPDH, PGK, BglK, & BglA-2) and their (co)substrates (FBP, NADH, Pi, ADP, MU-Glc).

| Concentration of LDH (μg/ml) | Z' | S/N |
|---|---|---|
| 5 | 0.91 | 550 |
| 2.5 | 0.84 | 500 |
| 1.25 | 0.83 | 360 |
| 0.625 | 0.78 | 220 |
| 0.313 | 0.73 | 140 |
| 0.156 | 0.57 | 78 |
| 0.0781 | 0.18 | 40 |
| 0.0391 | 0.32 | 20 |
| 0.0195 | -0.025 | 8.9 |
| 0.00977 | -0.50 | 4.9 |
| 0.00488 | -1.5 | 2.8 |
| 0.00244 | -5.4 | 1.0 |

**Supplementary Table S4.**

| Costs for standard assay by coupled enzyme cascade | | | | | |
|---|---|---|---|---|---|
| **Components Used** | **Supplier** | **Cost/unit ($CAD)** | **mg/unit** | **mg per rxn** | **Cost ($CAD)** |
| NADH | Sigma | 116 | 500 | 0.0035 | 0.000812 |
| MU-Glc | Sigma | 100 | 100 | 0.0169 | 0.0169 |
| G3P | Sigma | 407 | 8 | 0.00850 | 0.433 |
| GAPDH (GapA) enzyme | In-house | 250 | 10 | 0.001 | 0.025 |
| PGK enzyme | In-house | 250 | 10 | 0.001 | 0.025 |
| BglK enzyme | In-house | 250 | 10 | 0.001 | 0.025 |
| BglA-2 enzyme | In-house | 250 | 10 | 0.001 | 0.025 |
| | | | | | 0.550 |

| Costs for assay using extended coupled enzyme cascade | | | | | |
|---|---|---|---|---|---|
| **Components Used** | **Supplier** | **Cost/unit ($CAD)** | **mg/unit** | **mg per rxn** | **Cost ($CAD)** |
| NADH | Sigma | 116 | 500 | 0.0035 | 0.000812 |
| MU-Glc | Sigma | 100 | 100 | 0.0169 | 0.0169 |
| F6BP | Sigma | 129 | 1000 | 0.020303 | 0.00262 |
| FBA enzyme | In-house | 250 | 10 | 0.001 | 0.025 |
| TPI enzyme | In-house | 250 | 10 | 0.001 | 0.025 |
| GAPDH (GapA) enzyme | In-house | 250 | 10 | 0.001 | 0.025 |
| PGK enzyme | In-house | 250 | 10 | 0.001 | 0.025 |
| BglK enzyme | In-house | 250 | 10 | 0.001 | 0.025 |
| BglA-2 enzyme | In-house | 250 | 10 | 0.001 | 0.025 |
| | | | | | 0.170 |

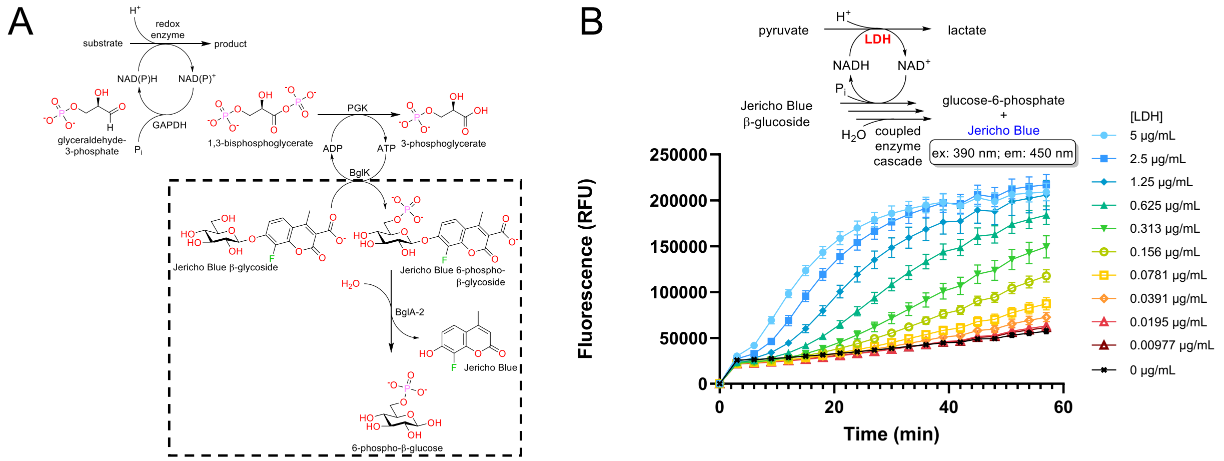


**Supplementary Figure S4.** (A) modified coupled enzyme cascade using Jericho Blue as a fluorescence reporter. JB-Glc was used in place of MU-Glc and changes from the standard coupled enzyme cascade are indicated by dashed line box. (B) Fluorescence-based assay of the reduction pyruvate catalyzed by LDH (at varied concentration) with a modified coupled enzyme cascade using JB-Glc (instead of MU-Glc).

**Supplementary Table S5**. Z' values and signal-to-noise ratio for LDH assays performed by fluorescence-based assay with a "Jericho Blue" reporter, using a coupled enzyme cascade including coupling enzymes (GAPDH, PGK, BglK, & BglA-2) and their (co)substrates (FBP, NADH, Pi, ADP, JB-Glc).

| Concentration of LDH (μg/ml) | Z' | S/N |
|---|---|---|
| 5 | 0.76 | 65 |
| 2.5 | 0.74 | 59 |
| 1.25 | 0.62 | 55 |
| 0.625 | 0.70 | 47 |
| 0.313 | 0.52 | 34 |
| 0.156 | 0.54 | 22 |
| 0.0781 | 0.053 | 11 |
| 0.0391 | -0.48 | 5.7 |
| 0.0195 | -2.2 | 2.78 |
| 0.00977 | -3.5 | 1.6 |
| 0.00488 | -20 | 0.36 |
| 0.00244 | -1800 | 0.0035 |

**Supplementary Table S6.** Plasmids used for expression of enzymes used in the coupled enzyme cascade(s)

| Plasmid | Protein encoded | Source |
|---|---|---|
| pTrcHis-BglK | BglK | [29] |
| pET28-SpBglA-2 | BglA-2 | [30] |
| pET28-gapA | GAPDH (*E. coli* GapA) | This study |
| pET28-pgk | PGK | This study |
| pET28-fbaA | FBA (*E. coli* FbaA) | This study |
| pET28-tpiA | TPI (*E. coli* TpiA) | This study |
| pET28-OYE2 | OYE2 | This study |

**Supplementary Table S7.** Primers used for cloning genes encoding enzymes expressed in this study.

| Target | Purpose | Sequence (5′ → 3′) |
|---|---|---|
| *gapA* | PCR for cloning (forward primer) | cgcgcggcagccatatgactatcaaagtaggtatcaacgg |
| | PCR for cloning (reverse primer) | gtcgacggagctcgaattcggatccttatttggagatgtgagcgatcag |
| | Mutagenesis (G188T+P189K; sense) | cagaaaaccgttgataccaagtctcacaaagactgg |
| | Mutagenesis (G188T+P189K; anti) | ccagtctttgtgagacttggtatcaacggttttctg |
| *pgk* | PCR for cloning (forward primer) | cgcgcggcagccatatgtctgtaattaagatgaccgatctg |
| | PCR for cloning (reverse primer) | gtcgacggagctcgaattcggatccttacttcttagcgcgctcttc |
| *fbaA* | PCR for cloning (forward primer) | cgcgcggcagccatatgtctaagatttttgatttcgtaaaacc |
| | PCR for cloning (reverse primer) | gtcgacggagctcgaattcggatccttacagaacgtcgatcgcgttc |
| *tpiA* | PCR for cloning (forward primer) | cgcgcggcagccatatgggtaactggaaactgaacg |
| | PCR for cloning (reverse primer) | gtcgacggagctcgaattcggatccttaagcctgtttagccgcttc |
| *OYE2* | PCR for cloning (forward primer) | tgccgcgcggcagccatatgccatttgttaaggactttaagc |
| | PCR for cloning (reverse primer) | cggagctcgaattcggatccttaatttttgtcccaaccgagtttt |
